# Distinct basal ganglia decision dynamics under conflict and uncertainty

**DOI:** 10.1101/2024.09.12.612658

**Authors:** Nadja R. Ging-Jehli, James F. Cavanagh, Minkyu Ahn, David J. Segar, Wael F. Asaad, Michael J. Frank

**Author notes:** School of Computer Science and Electrical Engineering, Handong Global University, Pohang-si, Republic of Korea (Minkyu Ahn). Department of Neurosurgery, UCSF Medical Center, San Francisco CA, USA. Co-senior authors (Wael F. Asaad, Michael J. Frank).

## Abstract

The basal ganglia (BG) play a key role in decision-making, preventing impulsive actions in some contexts while facilitating fast adaptations in others. The specific contributions of different BG structures to this nuanced behavior remain unclear, particularly under varying situations of noisy and conflicting information that necessitate ongoing adjustments in the balance between speed and accuracy. Theoretical accounts suggest that dynamic regulation of the amount of evidence required to commit to a decision (a dynamic “decision boundary”) may be necessary to meet these competing demands. Through the application of novel computational modeling tools in tandem with direct neural recordings from human BG areas, we find that neural dynamics in the theta band manifest as variations in a collapsing decision boundary as a function of conflict and uncertainty.

We collected intracranial recordings from patients diagnosed with either Parkinson’s disease (n=14) or dystonia (n=3) in the subthalamic nucleus (STN), globus pallidus internus (GPi), and externus (GPe) during their performance of a novel perceptual discrimination task in which we independently manipulated uncertainty and conflict. To formally characterize whether these task and neural components influenced decision dynamics, we leveraged modified diffusion decision models (DDMs). Behavioral choices and response time distributions were best characterized by a modified DDM in which the decision boundary collapsed over time, but where the onset and shape of this collapse varied with conflict. Moreover, theta dynamics in BG structures predicted the onset and shape of this collapse but differentially across task conditions. In STN, theta activity was related to a prolonged decision boundary (indexed by slower collapse and therefore more deliberate choices) during high-conflict situations. Conversely, rapid declines in GPe theta during low conflict conditions were related to rapidly collapsing boundaries and expedited choices, with additional complementary decision bound adjustments during high uncertainty situations. Finally, GPi theta effects were uniform across conditions, with increases in theta prolonging the collapse of decision bounds. Together, these findings provide a nuanced understanding of how our brain thwarts impulsive actions while nonetheless enabling behavioral adaptation amidst noisy and conflicting information.

## Introduction

We are constantly exposed to ambiguous and conflicting information, requiring us to carefully gather and assess information from various sources before making choices. When presented with conflicting information for alternative actions, it can be helpful to take time to pause and ensure that decisions appropriately reflect multiple sources of evidence. However, too much reflection can cause decision paralysis, especially amidst ambiguity. Therefore, striking a delicate balance, tailored to the circumstances, is crucial but also notoriously difficult. Here, we examine how such tradeoffs can be mitigated by dynamics in the basal ganglia (BG) that offer mechanisms to pause decisions and collect more evidence when needed but also to expedite choices when it may be come costly to accumulate for too long (1–5). We developed a novel perceptual paradigm which orthogonally varies conflict and uncertainty and examined how these conditions would lead to dynamic adjustments in decision strategies. Together with tailored computational models, we advance our understanding of the brain’s capacity to resolve trade-offs between impulsivity and behavioral adaptability with broad implications for biology, decision science, and health.

The BG comprise various subcortical structures that coordinates selection of actions in response to cortical inputs, while also regulating the needed evidence (i.e., decision threshold) for committing to a choice (1,6–9). Neurons in the striatum are the BG’s main input segment and accumulate evidence for alternative choice options (10,11). Those in the globus pallidus internus (GPi) are the BG’s main output segment and gate the striatal impact on decision-making (10–14). GPi receives input from two other BG structures, the globus pallidus externus (GPe) and the subthalamic nucleus (STN), which are part of distinct pathways that intricately link the BG and the cortex (7,8,15,16). To date, the relative contribution of these structures to decision-making, particularly in the presence of noisy and conflicting information, is not yet understood. We describe the distinct and complementary dynamics in the STN, GPe, and GPi for regulating decision-making within the same paradigm but orthogonally varying conflict and uncertainty.

Leveraging computational methods, we use diffusion decision models (DDMs; 28,29) to link activity in the various BG structures to specific aspects of decision-making dynamics (Fig. 1A). DDMs capture simultaneously *which* choices are made and *when* they occur across the full distribution of response times (RTs). They provide a powerful computational account for studying brain-behavior mappings because they decompose choices into distinct latent components that together resemble the dynamic decision process (19–21). Each component is represented by a quantifiable parameter with well-established psychological interpretation (19–21). However, the DDM is just one instance of a broader class of sequential sample models (SSM; 30), each with its own assumptions about the underlying decision dynamics (19,21–23). In this study, we show how one can leverage and test neurobiologically derived hypotheses for these alternative models of evidence accumulation.

**Fig. 1.**
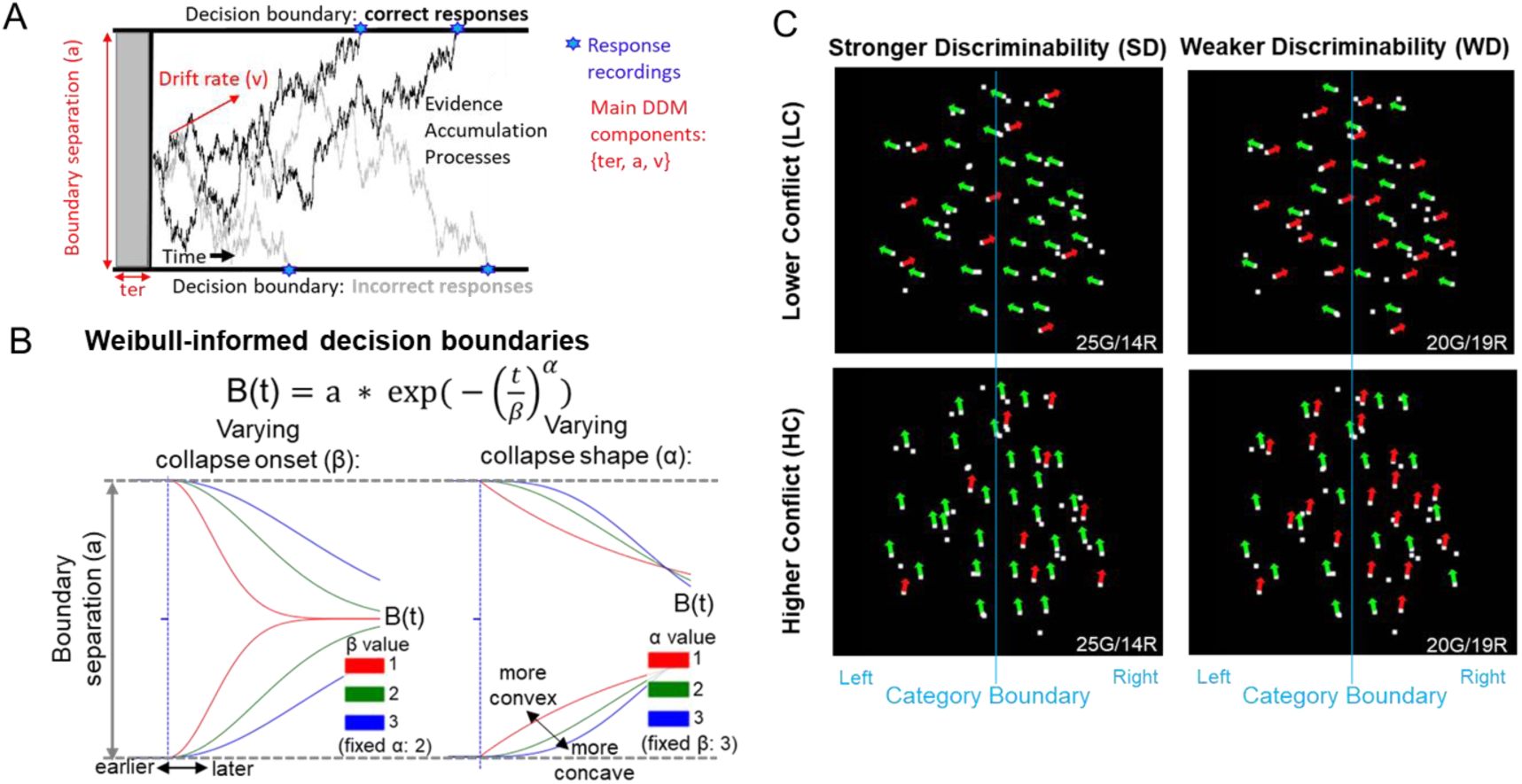
Characteristics of Weibull-informed decision boundaries index dynamic cautiousness in our experimental paradigm. **(A)** Representation of the classical diffusion decision model (DDM) with fixed decision boundaries. The DDM emulates latent decision-making processes that evolve over time and that represent the sequential accumulation of information. According to this model, choices, and corresponding response times (RTs) manifest from these decision-making processes. Specifically, decision-making processes are presumed to continue until they reach a certain decision threshold that is associated with a specific choice. **(B)** Dynamics of Weibull-informed decision boundaries, characterized by initial height *a* followed by a collapsing bound with separable onset (β) and shape (α) parameters. **(C)** Task stimuli involved dot motion patterns that varied in strong vs weak discriminability (motion coherence) and low vs high conflict (angle subtending response targets), producing four task conditions (SD-LC, WD-LC, SD-HC and WD-HC). For example, the SD-HC condition involved stronger discriminability but higher conflict because motion trajectories of dots were close to the category boundary (blue vertical line invisible to participants) of left/right responses (detailed in the Methods).

Inspired by past neurobiological models that focused on the neural dynamics across BG regions, we leveraged modified DDMs to advance and refine our current understanding of STN-mediated control over decisions. The STN, driven by cortical inputs, is known to detect *conflict* between choice alternatives (1,24–29) and pauses the selection of actions to promote further information integration or stops actions outright. The causal role of the STN as a *global brake* in response to decision conflict has been validated in behavioral, functional imaging, neural manipulation, and lesion studies across species (24,28,30–39). Past studies have used the standard DDM to show that elevations in cortical and STN theta band power, triggered by decision conflict, increases the DDM *decision threshold* parameter (also referred to as *boundary separation, a;* Fig. 1A), facilitating slower and more accurate responses (1,22,25,32,34,40,41). However, the standard DDM assumes that decision thresholds are constant during the decision-making process (Fig. 1A), whereas normative considerations such as those motivated above suggest that in many circumstances the boundaries should collapse over time (38,39,40,41, Fig. 1B). Moreover, neural data and network models suggest that STN and GP dynamics should translate into dynamic decision thresholds (22,25,28,37), thereby motivating modified DDMs with dynamic decision boundaries. We therefore tested whether the standard DDM or modified DDMs with dynamic boundaries best capture behavioral patterns in our paradigm. Applying dynamic DDMs in a mechanistic task that dissociates different forms of conflict and uncertainty allows us also to reconcile contradictory findings in previous studies. For example, many studies suggest that STN theta power increases decision threshold in the presence of higher conflict, but there are some indications that the same level of theta power actually reduces threshold under lower conflict (32,42). A collapsing decision boundary can help address this problem: instead of raising the boundary with conflict, one can start with a high boundary and merely prolong its collapse when there is conflict, but expedite its collapse when conflict is low so as to not waster further time accumulating evidence. Our findings confirm this hypothesis, which challenges a simple unidirectional association of higher STN theta power with increased decision threshold (28,43,44). Moreover, we reveal a prominent role for reductions in GPe theta under low conflict situations in which decisions can be expedited.

Past studies vary in terms of what constitutes *conflict*, from perceptual uncertainty, to uncertainty in value-based decision-making, to response or stimulus conflict (24,32,36,42,45). Dissociations between uncertainty and conflict within the same paradigm are missing. We will show that this dissociation reveals novel mechanisms of how BG structures contribute to decision dynamics, potentially involving interactions between the STN, GPe and GPi (8,24,32). Neurocomputational simulations suggest that the GPe, via strong reciprocal interactions with the STN, could play pivotal roles under higher uncertainty by expanding the potential spectrum of decision boundary dynamics (1,46). The GPe’s promotion of response cautiousness may be particular effective for resolving conflict under conditions of high information uncertainty. This might be because the GPe has been tied to fine-tuning processes for selective attention and has shown greater specificity in information transmission than other BG output structures (1,7,8,47,48).

To examine how distinct BG components contribute to decision dynamics, we recorded local field potentials (LFPs) from STN, GPi, and GPe in human patients with Parkinson’s disease (n=14) or dystonia (n=3) while they judged the primary direction of moving dots, either left or right (Fig. 1C). In this task, we independently varied conflict and uncertainty in discriminability (detailed in the Methods and below). We then related variations in single-trial theta band dynamics to the dynamic decision boundaries in the modified DDM. We focused on theta power based on strong a priori hypotheses established by previous work (34,42,49) and to focus our main findings on the novel dissection of decision dynamics across three distinct BG structures. Importantly, we also confirmed that a variant of this task (see Methods) adapted for younger students without neurological disorders (n=25) produced similar conflict-induced behavioral changes as those found in the patient groups, establishing that the behavioral dynamics are generalizable.

As hypothesized based on past neurocomputational applications (22,28), the modified DDM with dynamically collapsing decision boundaries best captured behavioral patterns. The onset and shape of this within-trial dynamic decision threshold was characterized by a Weibull distribution governed by two free parameters (Fig. 1B). Early (post-stimulus) theta activity modulated collapse onset, whereas later pre-response activity modulated collapse shape.

Theta dynamics in distinct BG regions modulated the decision boundary collapses in a complementary fashion. To preview our main findings: Under strong motion coherence, the presence of conflict increased STN theta and prolonged collapsing boundary, supporting more cautious and accurate choices. In contrast, under low conflict, decisions could be made expeditiously, and we found that decreased GPe theta in this case promoted a more precipitous decline in the collapsing boundary. When motion coherence was weak (higher uncertainty), boundary collapse was delayed with higher theta in both STN and GPe. Finally, higher GPi theta was related to prolonged decision boundaries uniformly across task conditions, consistent with its role as the final output structure of the BG (1,27).

## Results

### Separating conflict from discriminability

In our perceptual decision task, we independently manipulated uncertainty in discriminability (coherence) and conflict (Fig. 1C). Specifically, task stimuli involved moving dot patterns that varied in two levels of motion coherence (signal strength) and angular trajectory (signal interference), respectively. Varying motion coherence makes perceptual discriminability stronger or weaker according to the degree of overlap of cortical populations coding for specific motion directions (8,20,50–55). Varying angular trajectory induces cognitive conflict: even when coherence is high, those trajectories close to the category boundary (vertical blue lines in Fig. 1C) induce conflict due to the overlap of category-specific cortical populations (56–58). Since priming bimanual responses with the requirement of a single motor output has been advanced as a formal definition of cognitive conflict (59,60), we refer to these conditions as lower vs. higher conflict (see Methods). Within the SSM framework, discriminability affects the rate of sensory evidence accumulation, whereas co-activation of mutually incompatible responses elevates decision boundaries (22,34,42,59).

As typically seen in dot motion discrimination tasks, stronger compared to weaker discriminability was associated with higher accuracy (mean difference: 8.32%, SEM = 1.74%, p < 0.001)^1^ and faster mean RTs for correct responses (mean difference: -100ms, SEM = 28ms, p = 0.004).^2^ We augment past findings by showing that higher conflict induced slower mean RTs for correct but not for incorrect responses, and more so under stronger than weaker discriminability (Fig. 2A). The absence of conflict-induced slowing for error responses can be linked to failures to sufficiently increase decision thresholds (29,30). Specifically, higher accuracy is associated with increased RTs (reflecting a so-called speed-accuracy trade-off up to a saturation point) as shown in Fig. 2B. We provide additional summary statistics in Supplementary Fig. 1, aside from the empirical RT quantiles and accuracy by task conditions (Fig. 2D).

**Fig. 2.**
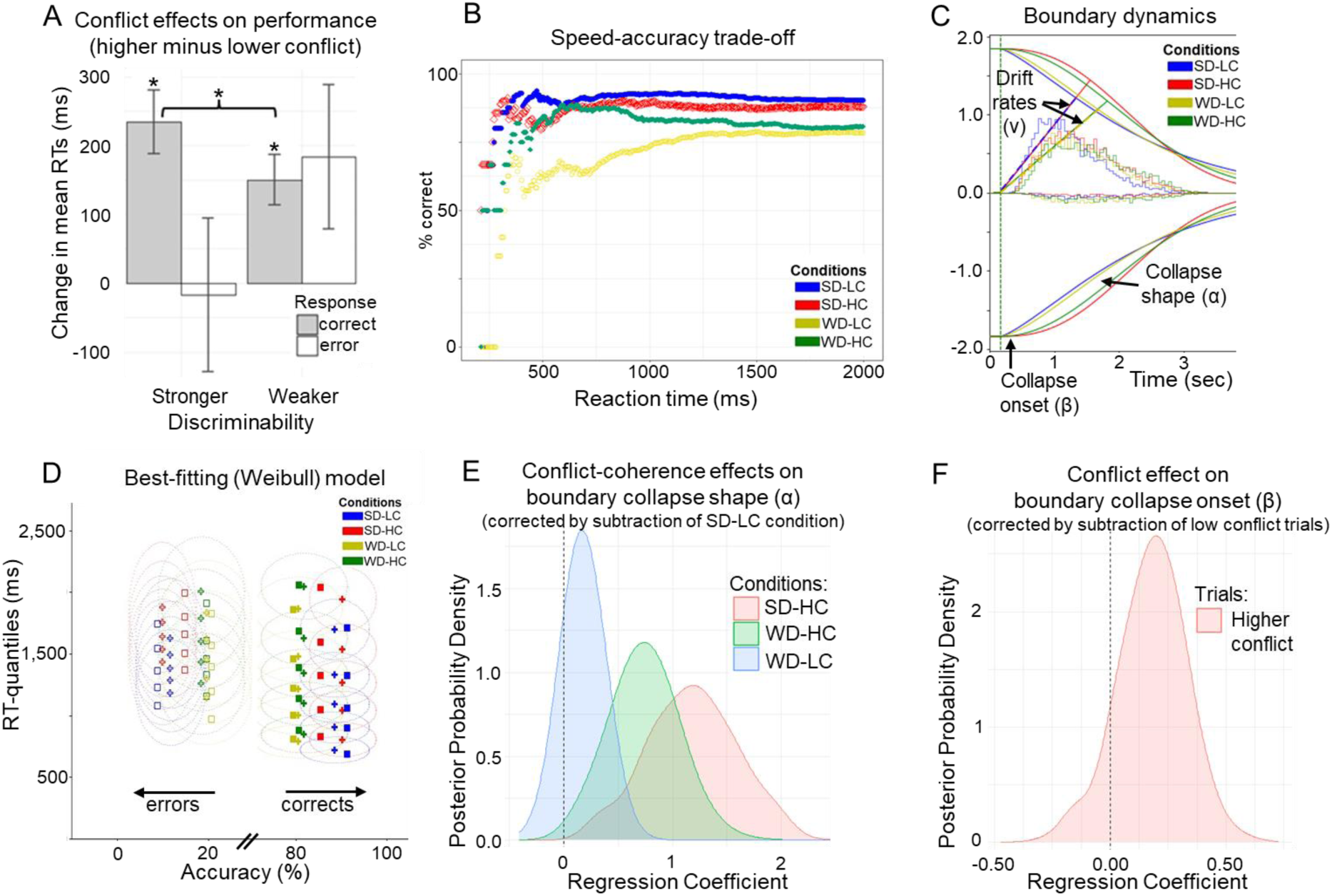
Behavioral task performance. **(A)** Changes in performance measures (high minus low conflict) due to conflict for each discrimination level (stronger = easier, weaker = harder). Error bars indicate SEMs. Asterisks indicate significance (p<0.05; Wilcoxon signed-rank tests for paired samples). **(B)** Accuracy (% correct) as a function of reaction times with shaded intervals indicating SDs. Task conditions: SD-LC=stronger discriminability, lower conflict; SD-HC=stronger discriminability, higher conflict; WD-LC=weaker discriminability, lower conflict; WD-HC=weaker discriminability, higher conflict. **(C)** Best-fitting dynamics of boundary collapse and drift rates by condition using the Weibull model. Whereas stronger discriminability increases drift rates, higher conflict induces a more prolonged elevation in decision bound before collapsing. **(D)** Posterior predictive check of best-fitting model. Squares indicate data; crosses indicate posterior predictions. Ellipses surrounding crosses indicate 95% confidence intervals in expected range of data given stochasticity in model and estimation uncertainty. All measures were calculated by condition and by subject before averaging. **(E)** Conflict-by-coherence interaction, leading to more concave boundary collapse for high conflict particularly under stronger coherence (SD-HC). Shown are differences in posterior distribution of collapse shapes (α) of conditions relative to the easiest SD-LC condition. Posterior probability (PP): α_SD-HC_ > α_SD-LC_ = 1.00; α_WD-HC_ > α_SD-LC_ = 0.9973; α_WD-LC_ > α_SD-LC_ = 0.8882. **(F)** Main effect of high-low conflict on the onset of boundary collapse. Shown is the difference in posterior distribution of collapse onsets (β) between higher versus lower conflict trials. Posterior probability (PP): β_higher_ > β_lower_ = 0.8937.

### Evidence for dynamic cautiousness adaptations during decisions

We fit a variety of SSMs that varied in either decision boundary dynamics (i.e., fixed, linear collapsing, Weibull-informed collapsing) or relative (stochastic) evidence accumulation dynamics (i.e., constant versus variable drift rates). Supplementary Table 1 provides model comparison and posterior predictive checks. Supporting the central behavioral prediction, the best-fitting model included constant drift rates that varied by discriminability (stronger, weaker), and Weibull-informed collapsing decision boundaries whose onset and shape varied by conflict and discriminability-by-conflict interaction, respectively. Other model parameters were fixed across task conditions (see Methods). This best-fitting model produced dynamics are graphically simulated in Fig. 2C, showing both drift rate and boundary effects as a function of discriminability and conflict. It demonstrated good parameter recovery (Supplementary Fig. 2) and captured the data well (Fig. 2D: squares representing empirical data are within ellipses representing Bayesian estimation uncertainty). It particularly captured the tails of the RT distributions (including accuracies) better than the full DDM with variability parameters (Supplementary Table 2). Importantly, the students’ behavioral pattern was also best fit by the Weibull DDM, with similar modulations of drift rates by discriminability and decision boundary dynamics by conflict and discriminability (Supplementary Table 3). This suggests that the presented patterns for the patients are generalizable.

### Distinct cautiousness dynamics tied to conflict and discriminability

Using Bayesian hierarchical model estimations, we report parameter changes across task conditions in terms of their corresponding posterior *probabilities* (PP) that index the likelihood of observed effects exceeding zero (61,62). Drift rates (v) were larger under stronger than weaker coherence discriminability (PP: v_stronger_ > v_weaker_ = 1.00), consistent with past studies that manipulated discriminability without independently varying perceptual conflict (20,32,63). Moreover, conflict induced a prolongation in the boundary collapse, an effect that is captured by a combination of the collapse onset and its shape. The shape parameter (α) was more concave for higher than lower conflict trials (Fig. 2E; PP: α_SD-HC_ > α_SD-LC_ = 1.00; α_WD-HC_ > α_WD-LC_ = 0.99); and more so under stronger than weaker discriminability (PP: α_SD-HC_ > α_WD- HC_ = 0.93). Collapse onset (β) was marginally delayed for higher than lower conflict trials (Fig. 2F; PP: β_higher_ > β_lower_ = 0.89).^3^ Notably, we show below that this relationship was moderated by the magnitude of theta activation and its distinct effects across BG components.

### Conflict-related theta dynamics across all BG components

Previous studies have established that cognitive conflict increases theta band activity in the STN (26,29,36,37,42,49). Fig. 3A (left panel) shows greater post-stimulus theta increases in STN and GPe for higher than lower conflict trials. This conflict-related theta activity persisted in the STN leading up to the response, before finally declining, for both discriminability levels (Fig. 3A: right panel). This pattern might be expected if STN theta prolongs the bound under high conflict before collapsing, a claim we will test formally below. In contrast, while GPe theta band dynamics were high for both high conflict conditions, they showed an early and rapid decline in the high coherence case (the condition in which one should not need to accumulate any more evidence because conflict is low and discriminability is high). We will also test this effect on boundary collapse below. During the pre-response period, both the GPi and STN showed comparable theta responses, supporting the notion that these components act in synchrony (7,8,64). We provide additional time-frequency plots in the Supplementary Fig. 5-9, demonstrating that in addition to theta, STN showed specific decreases in beta-frequency power (13–30Hz) leading up to the response, consistent with past research showing beta desynchronization in STN prior to motor engagement (36,65–67). Fig. 3B summarizes the time-frequency plots of conflict-related changes across BG components. We provide additional time-frequency plots for each discriminability level and each task condition (Supplementary Fig. 5 & 6). Moreover, supplementary Fig. 7-9 provide event-related potential (ERP) analyses and phase-locking value (PLV) analyses demonstrating the reliability and specificity of neural responses during task performance to confirm that the observed neural activity is indeed tightly linked to relevant behavioral processes.

**Fig. 3.**
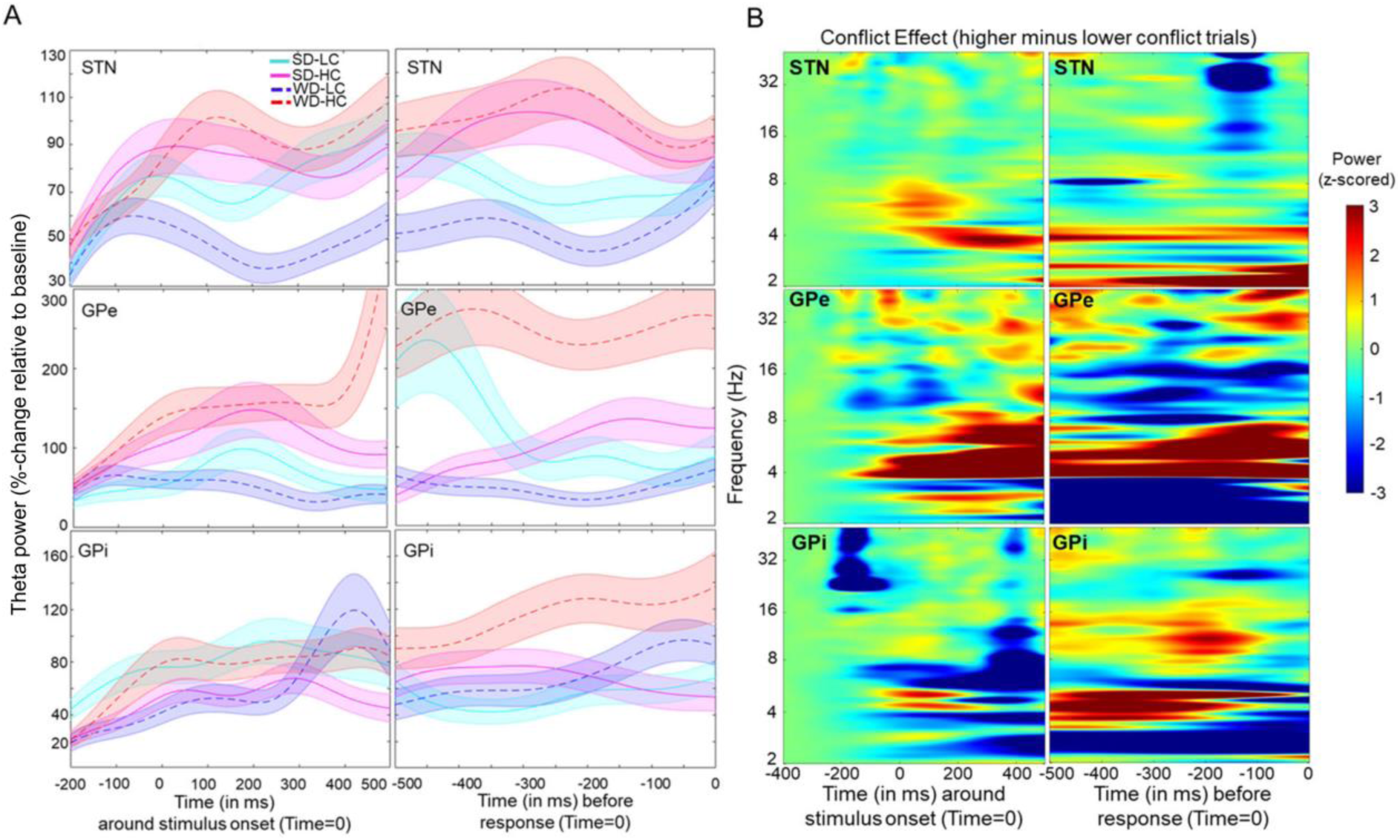
Task-evoked neuronal response. **(A)** Stimulus-induced mean changes (solid/dotted bold lines) in theta power (4–8Hz) by condition. The time series of each trial was normalized and averaged across channels (for each BG component). Shaded areas represent within-subject standard errors. Task conditions: SD-LC=stronger discriminability, lower conflict; SD- HC=stronger discriminability, higher conflict; WD-LC=weaker discriminability, lower conflict; WD-HC=weaker discriminability, higher conflict. **(B)** Time frequency plots show a task-evoked increase in LFO power (averaged across channels) relative to baseline. Spectra are shown for high minus low conflict (averaged across coherence) aligned to stimulus onset (left panel) and response (right panel) for each BG component.

### Theta modulates dynamic cautiousness with collapsing boundaries

We next assessed the functional significance of these neural dynamics with the modified (Weibull) DDM. To do so, we added trial by trial measures of theta activity as neural regressors into the model discussed above. We found that model fits improved over those applied to behavior alone. Specifically, including early (post-stimulus) theta activity (Fig. 4A) and later (pre-response) theta activity reduced the DIC values by 181, suggesting that neural measures governed the decision boundary dynamics on a trial-by-trial basis. Specifically, post-stimulus theta activity modulated collapse onset (β), while later pre-response theta activity modulated collapse shape (α). To recapitulate, we found increased theta activity in response to conflict across all BG components (previous section). In this section, we showed that trial-wise markers of these conflict signals are predictive of the dynamics of collapsing boundaries. Next, we show that the impact of these dynamics on collapsing boundaries differed by region and by task condition, in line with differential mechanisms needed to prolong or rapidly collapse the boundary as a function of conflict and uncertainty.

**Fig. 4.**
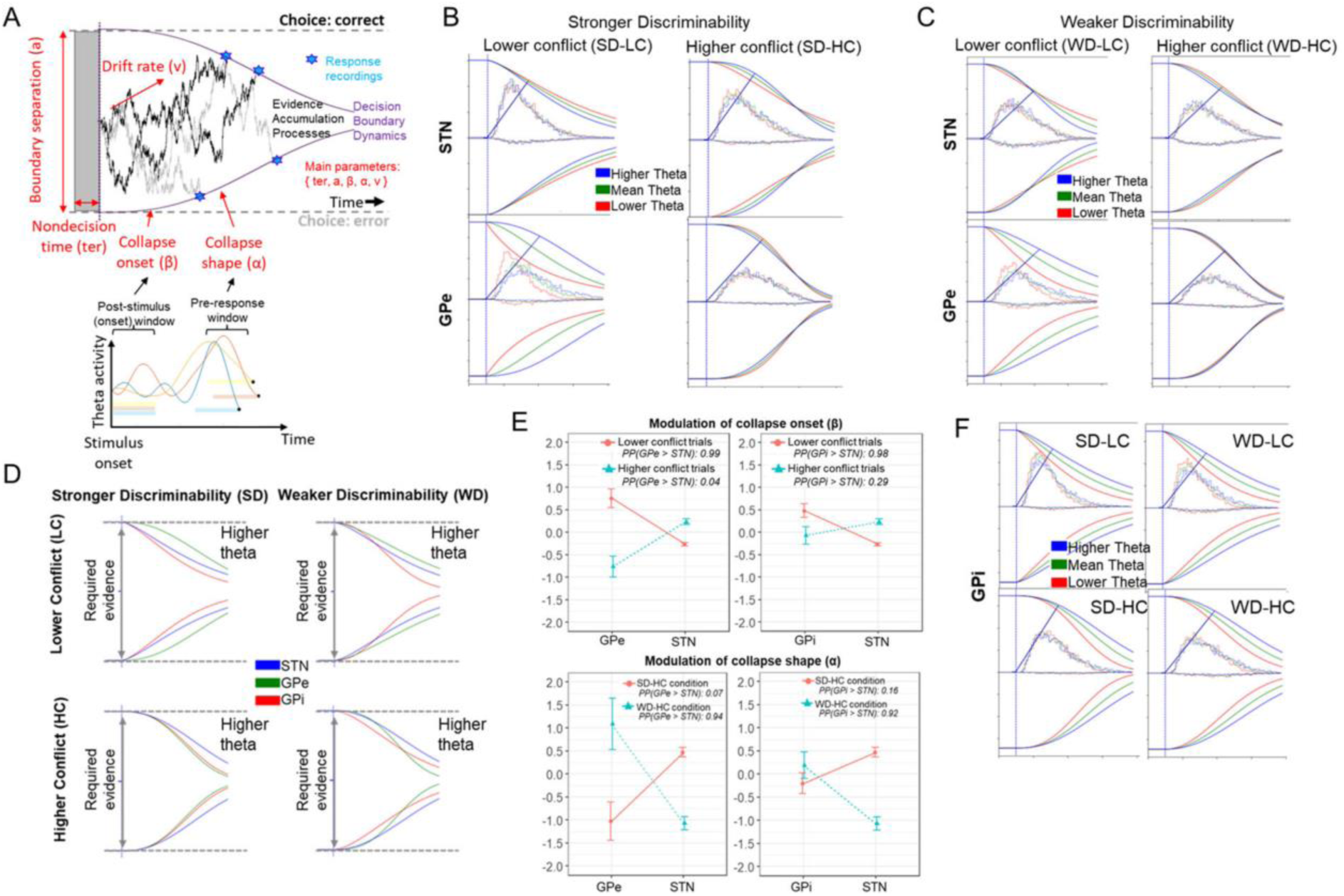
Decision boundary dynamics modulated by theta activity. **(A)** Exemplified scheme for integrating mean activation in theta-frequency band during pre-response (z-scored mean activation during the last 500ms before response) and post-stimulus (z-scored mean activation during the first 500ms after stimulus onset) periods as trial-based neural regressors into the best-fitting model. Single-trial LFPs were z-scored for each condition separately before entering them into the HDDM. Means refer to hierarchically centered grand means. **(B)** Theta-specific modulations for STN and GPe under stronger discriminability. STN theta was related to prolonged decision boundary in SD-HC but decreased boundary in lower conflict conditions. Conversely GPe theta was related to a prolonged bound in low conflict. **(C)** Theta-specific modulations for STN and GPe under weaker discriminability. **(D)** Theta-specific modulations across all BG components for higher theta activity. **(E)** Comparison of collapse onset (β) and collapse shape (α) across BG components for higher theta (i.e., theta power 1SD above hierarchically centered grand means). Coefficients refer to the group posterior distributions (whereby points refer to means and vertical lines refer to SEMs). We also present the posterior probability (PP) for group-specific differences in coefficients below each plot. See table S3 for all estimated (posterior) coefficients. **(F)** GPi theta was related to prolonged bound across all conditions. For visualization, we show impact of theta at three levels (i.e., 1 SD below grand mean, at grand mean, 1 SD above grand mean; but all regressions were done continuously).

### STN and GPe differentially prolong and collapse boundaries in high vs low conflict

Previous findings demonstrated that conflict increased frontal and STN theta, which elevated boundary separation, facilitating response caution (32,34,35,42,68,69). Consistent with past findings, under stronger discriminability conditions, we found that increased STN theta was linked to elevated decision boundary on higher conflict trials (PP: α_SD-HC,θz|higher_ > α_SD-HC,θz|mean_ = 0.93). Compared to past findings though, we show that boundary is not statically elevated at trial onset but instead that conflict induces more concave collapse of the boundary rather than a static (i.e., a priori and constant) elevation of the boundary, and that this collapse shape was governed by STN activity leading up to the behavioral response (Fig 4B). This effect was not observed in low conflict trials; instead, higher STN theta was linked to somewhat more rapid onsets of boundary collapses under lower conflict (PP: β_θz|higher_ < β_θz|mean_ = 0.97). This provides empirical evidence for hypotheses raised by previous studies (32,42). In summary, increased theta activity in the STN modulated decision boundaries to preferentially collapse more slowly during high-conflict situations but sped up this collapse during low-conflict situations.

Strikingly, we observed diametrically opposed effects in GPe, which were related to much more rapid collapses of boundary adjustments in lower conflict trials (Fig 4B, bottom row), consistent with the rapid theta decline during the pre-response period shown in Fig. 3A (middle-right panel). Specifically, earlier (post-stimulus) decreases in theta were related to a more rapid collapse onset (β_θz|higher_ > β_θz|mean_ = 0.96) in lower conflict trials^4^. Hence, decreasing GPe theta expedites the inevitable collapse. In summary, increased STN theta delayed boundary collapses, leading to more cautious and accurate decisions under higher conflict, while decreased GPe theta fastened boundary collapses under lower conflict. Moreover, under stronger discriminability, higher theta in the STN was associated with more concave collapse shape, while the opposite was found in the GPe.

### Complementary dynamics in STN and GPe for weaker discriminability

In the weaker discriminability condition, STN modulation of boundary collapse was significantly different from that reported above for stronger discriminability (Fig 4C: α_SD- HC,θz|higher_ > α_WD-HC,θz|higher_ = 0.95). These results converge with with biophysical models showing that STN theta requires strong cortical inputs across two conflicting responses, which have supralinear effects on theta activity (28); see also (1). In other words, the STN seems to exhibit less pronounced engagement in boundary regulation under weaker discriminability in which conflict might be less salient than under stronger discriminability.

For GPe, unlike the strong discriminability case, theta power did not decline rapidly in weak discrimability (Fig 3). Instead, higher theta related to more concave collapses on higher conflict trials, promoting more cautious responding (PP: α_WD-HC,θz|higher_ > α_WD-LC,θz|mean_ = 0.97). This suggests that after the early collapse onset, the pre-response decision period is still modulated by theta power: continuing to buy time in GPe during weaker evidence (WD-HC condition). In summary, conflict-related heightened theta activity in the GPe later in the decision-making process modulated boundaries to collapse more slowly during harder decisions (involving weaker discriminability) but sped up this collapse during easier decisions (involving stronger discriminability and lower conflict). This shows the complementary dynamics of STN and GPe depending on discriminability levels (Fig. 4E).

### Universal dynamics in GPi for all discriminability levels

Higher theta in GPi was uniformly linked to delays in the onset of the collapse rather than its shape, and these effects were consistent across task conditions (Fig 4F). These findings are consistent with the notion that GPi neural activity governs BG output to coordinate action selections in a task-independent fashion, whereas opponent STN and GPe signals have different effects on GPi depending on the decision-relevant factors. Overall, trial-by-trial modulation in post-stimulus theta activation modulated response cautiousness by varying collapse onset (but not shape) in distinct ways across the BG components (see Supplementary Table 4 and Fig. 10).

## Discussion

The basal ganglia (BG) play a pivotal role in decision-making processes across various species, yet the field lacks comprehensive data, particularly in humans, on how neural dynamics within different BG structures facilitate these processes. Especially, the dynamic interplay between these mechanisms and their contextual adaptation amidst noisy and conflicting information, crucial for preventing impulsive actions and fostering adaptability, remains poorly understood. Our study uniquely investigates the involvement of decision dynamics across multiple BG structures in humans. This is an overlooked perspective as past research has predominantly concentrated on the subthalamic nucleus (STN) or the globus pallidus internus (GPi) in isolation, and not how either of these structures’ ongoing activity contribute to decision dynamics in the form of collapsing bounds. Moreover, unlike past studies, we distinctively separate the effects of conflict from other forms of decision uncertainty. Our findings underscore the significance of understanding different mechanisms for controlled decision-making, providing insights into the relevant dynamics of impulsivity.

Our study defined specific computational roles for population-level neural activities across BG components that underlie decision-making. Recent studies with rodents and monkeys have identified that the BG causally contribute to the accumulation of sensory evidence over time (10,11,70). These observations are consistent with computational models of BG circuitry in which BG output “gates” the selection of cortical actions with a dynamic threshold determining the amount of evidence needed to commit to a choice (1,22). Our observations characterize the neural implementation of these algorithmically defined processes across three BG structures.

Our study presents novel decision dynamics within the basal ganglia, utilizing unique intracranial data and leveraging novel computational methods to rigorously assess the links between neural and decision dynamics within a novel conflict paradigm. Previous research in this area has assumed a static decision boundary, largely due to technical limitations in model estimation, which are now overcome via our novel model methods. The collapsing boundary model is debated in many aspects of decision making largely based on behavioral data (e.g., (71) but see: (72,73)) and our study provides the first characterization of how it can be implemented by subcortical dynamics as a function of conflict and uncertainty. We found that SSMs with dynamic decision boundaries captured behavioral patterns better than classical DDMs with time-invariant decision thresholds (2,5). Such a dynamic process is also justified by underlying neural dynamics and by normative considerations (2,3), especially when task demands involve a mixture of difficulty levels across trials (4,5). By quantifying these boundary dynamics, we further demonstrated that theta activities in the STN do not uniformly raise decision boundaries. Instead, they modulate boundary collapse over time, with opposing effects in higher versus lower conflict. While previous biophysical model simulations have suggested these opposing effects (28), our study is the first to empirically demonstrate them and link them to decision-relevant boundary collapse by utilizing modified DDMs.

Many decision-relevant dynamics in the STN and the GPe complemented each other. First, whereas STN theta was largely related to boundary adjustments in higher conflict trials, GPe theta modulation was strongly related to a rapid collapse of the decision bound in lower conflict trials. Moreover, these effects were especially prevalent in STN under stronger discriminability but by the GPe under weaker discriminability. This latter finding is consistent with the notion that the GPe may serve to guide selection of specific actions rather than exert global braking, aligning with the selective versus global model commonly associated with the indirect and hyper-direct pathways (25,48,49,64). Overall, the modulatory roles of these hyper-direct and indirect pathway structures differed from GPi dynamics, which uniformly related to prolonged decision boundaries across task conditions, supporting the notion that this structure forms the final stage of BG output that is subsequently used for coordinated action selections, and consistent with related findings in monkeys (3).

We leveraged the unique opportunity of subcortical neural recordings in patients with PD or dystonia to dissect decision-relevant dynamics across different BG structures. This special population limits the potential generalizability of these findings, although there is no reason to think that these subcortical operations are different in those without neurological disorders (48,65,66). Indeed, comparison to a group of college students without neurological conditions suggests that behavioral patterns and model dynamics were similar across groups. We acknowledge that the uneven distribution of diagnoses and the small sample size (Supplementary Table 1) may limit the generalizability of our results. Future studies are necessary to validate and extend our findings. Additionally, subsequent studies could explore similarities and differences between Parkinson’s disease (74,75) and Dystonia (76). While our findings do not address potential differences between these clinical conditions, Supplementary Fig.11-14 might provide insights for future research.

While we extracted decision-relevant neural dynamics in the theta frequency band (4–8Hz), previous studies also suggest decision-relevant dynamics exist in the beta frequency band. However, beta-specific dynamics seem particularly important for stopping prepotent actions (33,77,78) or resolving unambiguous decision conflict (36,79). Understanding the extent to which these findings generalize to other dynamic decision tasks, and the possible role of activity in other frequency bands, will require further investigation. Our study focused on the role of theta dynamics in decision-making processes across multiple BG structures by utilizing specially modified DDMs to separately examine the dynamic effects of conflict and uncertainty. This approach provided evidence that distinct decision dynamics are linked with different slowing mechanisms.

Cognitive processes related to information integration and choice initiation are often formalized using SSMs, with the DDM being the most prominent application (30). In our study, we demonstrate that an adaptation of the DDM, incorporating more biologically plausible collapsing decision boundaries, best represents behavioral patterns in a perceptual decision task with varying levels of both conflict and uncertainty. Additionally, neural signals in specific interconnected regions of the basal ganglia—a subcortical network linked to the frontal cortex’s learning and planning systems— modulate key parameters of this modified DDM complementarily and on a trial-by-trial basis. Specifically, GPe shortened decision processes during lower conflict yet prolonged them under higher uncertainty, whereas STN operated in higher conflict and lower uncertainty scenarios. GPi effects were uniform across conditions.

Our findings may be clinically useful because they suggest new possibilities to better understand multi-faceted symptoms like impulsivity, not only for Parkinson’s disease, but also for other conditions such as attention-deficit hyperactivity disorder (19,80–82). Furthermore, more broadly, our model with dynamic decision boundaries can be used to distinguish among various control processes known to influence decision-making at different time points. For instance, early- and late-stage control processes are often differentiated in task-switch paradigms and research indicates that distinguishing between these different control processes enhances our understanding of the impact of aging on cognitive control (83–85). Moreover, our findings demonstrate that varied theta dynamics correlate with unique control mechanisms within different BG components, and the interplay of these processes makes us both deliberate in our actions and capable of adapting to change. As the prominence of neuromodulation and neurofeedback continues to rise, understanding how to target and regulate anatomically and functionally distinct neural mechanisms, such as these, becomes increasingly crucial.

## Data availability

Data will be available with manuscript publication.

## Materials and methods

### Participants

Participants included N=17 patients with either Parkinson’s disease (PD; n=14) or dystonia (n=3) who were undergoing implantation of deep brain stimulation electrodes. The decision to undergo routine, awake surgery was made by a multidisciplinary clinical team without any consideration of research related factors. All participants provided informed consent prior to surgery, and the Institutional Review Board of Lifespan / Rhode Island Hospital approved the study. All study activities were carried out in accordance with the principles outlined in the Declaration of Helsinki.

All recordings were performed in the dopaminergic OFF state as is standard for awake Deep Brain Stimulation (DBS) procedures. Electrode implantation targeted either STN or GP (i.e., the latter targeting either the internal or external segment). Many participants completed multiple sessions with recordings from different locations in STN, GPe and/or GPi. Out of 40 recording sessions in 17 patients, three sessions were excluded due to corrupted data (one patient), seven for fewer than 40 trials completed (six patients) and four for chance performance on the task defined as <50% total accuracy (two patients). From the remaining 26 sessions, 15 STN recordings were from eight patients, and 11 GP recordings (five GPe, six GPi) were from six patients. Recordings were from the left side in 23 of the 26 sessions, and 14 out of 16 of these patients used their right hand for the task. A total of seven sessions included recordings from patients diagnosed with Dystonia (four GPe, three GPi). Supplementary Table 1 provides details about the diagnosis, handedness, and other relevant information for each subject.

We also collected data from a group of undergraduate students (N=25) without any diagnosed neurologic conditions. This allowed us to test whether behavioral patterns across task conditions were specific to the patient groups or also observable in those without neurologic illness. This study was approved by the University of New Mexico (UNM) Institutional Review Board and all participants provided written informed consent. Participants received course credit for participation and the average age was 19.4 years old (SD=1.3).

### Cognitive task

All participants completed a varied number of trials involving a moving dots kinetogram programmed in MonkeyLogic (86). Each trial consisted of 100 white dots (three pixels) on a black background moving in a circular aperture (Fig. 1C). All dots had at least three frames of consecutive movement and each dot was replaced in a proportional stepwise fashion. The task design is similar to a previous animal study (58). For this study, 50% of dots always moved in random vectors. The remaining 50% of dots were split between leftward and rightward directions of movement. The subsequent task specifics differed slightly between the patient and student groups to avoid any ceiling effects in behavioral measures.

For the patient groups, 36% of dots moved towards the target direction on stronger trials, while 14% of dots moved towards the other target. On weaker trials, 30% of dots moved towards the target while 20% moved in the opposite direction. Extensive pilot testing revealed that these countervailing directions yielded maximally dissociable outcomes, particularly given the additional manipulation of angular direction. In the context of sequential sampling models, this change in dot coherence was expected to alter the drift rate of evidence accumulation. Moreover, we aimed to alter decision threshold with a manipulation of dot angular trajectory. The 50% of non-randomly moving dots either moved in oblique left or right angles (112 or 248 degrees) or in tightly vertical left or right angles (170 or 190 degrees). This manipulation was specifically designed to prime unidirectional vs. bidirectional responses. Since priming bidirectional responses with the requirement of a single motor output has been advanced as a formal definition of cognitive conflict (59,60), we refer to these conditions as lower vs. higher conflict. In sum, the experiment consisted of a cross-over two (coherence: stronger, weaker) by two (conflict: lower, higher) manipulation designed to alter drift rate and decision threshold respectively.

Binary choices in such perceptual tasks depend on judgments made relative to a decision criterion (shown as blue vertical lines in Figure 1C), which differentiates left from right responses. The proximity of the individual dot motion trajectories to this criterion determines the level of conflict: dots moving on a more acute angle (relative to the vertical decision criterion) create more conflict (due to activation of multiple category-specific cortical populations) than those clearly aligned with a specific right or left response option (56–58). In sum, the trajectory angle of the dots determines higher or lower conflict levels, while the dot coherence determines stronger or weaker discriminability, indicating the relative strength of evidence for left versus right responses.

The task specifics were slightly adjusted for the student group. Specifically, while coherence level on easier trials (i.e., stronger discriminability) was set to 36% for the patient groups, this value was decreased to 34% for the students. Moreover, the angle difference for higher and lower conflict trials was set to 20 and 126 degrees, respectively, for the patient groups. These differences were adjusted to 60 and 120 degrees, respectively, for the students. All other task specifics were the same across all participants. See Supplement (Section 1) for information on response devices.

### Electrophysiology

DBS targeting was performed using a combination of indirect (AC-PC coordinate system), direct (MRI target visualization) and neurophysiological methods (see Supplement, Section 1).

Electrophysiology was recorded from patients using clinical microelectrodes sampled at either 40k or 44k Hz using the AO FDA-approved human neurophysiology system and downsampled to 1000 Hz for local field potential (LFP) processing. Three to four signal channels were simultaneously acquired in any given task session, and analyses were performed on the average post-processed LFPs of simultaneous signals from the same brain structure. LFPs were time locked to the stimulus onset in -2000 ms to 5000 ms epochs; these were then shifted by the RT to derive response-locked LFPs. LFPs were low pass filtered at 20Hz and baseline corrected to the time locking event (defined as t = 0 ms).

Power was normalized by conversion to a decibel (dB) scale (10xlog10(power/ power baseline)), allowing a direct comparison of effects across frequency bands. The baseline for each frequency consisted of cross-condition averaged power from -500 to -300 ms prior to the onset of the trial. Analyses of theta-band (4–8 Hz) power used a Hilbert transform of band-pass filtered data. For HDDM single trial regression analysis, epochs were rejected if the theta filtered power envelope exceeded three standard deviations from the mean.

For theta-specific frequency plots, we averaged the theta-band time series (based on Hilbert transformation) across trials (for a given condition) to show the percentage change in power. To do so, we normalized the theta-band time series by subtracting the mean of the trial-specific baseline period (from -500 to -300ms prior to stimulus onset) to show the respective within-subject standard errors. Each epoch was then extracted (for stimulus onset: -400 to +500 ms whereby 0 reflects stimulus onset; pre-response: -500 to 0ms whereby 0 reflects response). For the time-frequency plots, we computed the LFPs using the continuous wavelet transform with a width (i.e., cycles) of each frequency band set to four plus f divided by 6 where f refers to frequency. We then standardized power of each frequency by subtracting the trial-specific baseline period (from -500 to -300ms prior to stimulus onset) from the time series of that frequency and dividing by the standard deviation of that baseline period. Trials with excessive activity during the baseline period were excluded. Each epoch was then extracted (for stimulus onset: -400 to +500 ms whereby zero reflects stimulus onset; pre-response: -500 to 0ms whereby 0 reflects response).

### Summary statistics of behavior

Due to the low sample size, we used non-parametric tests for analyses of summary statistics such as mean reaction times (RTs) and accuracy. We first tested the effect of discriminability (stronger versus weaker) on mean RTs for correct and error responses to validate that our coherence manipulations had intended behavioral effects that were typically seen in dot motion discrimination tasks. We then examined conflict-induced effects on mean and quantile RTs (for corrects and errors) and accuracy for each discriminability level. Results from these analyses are reported in Fig. 2 and in Supplementary Fig. 1. All statistical comparisons are based on paired Wilcoxon signed rank tests in R (Version 4.1.2; 74) with an alpha of 0.05.

### Sequential sampling modeling

We used the new LAN (Likelihood Approximation Networks) extension of the HDDM toolbox that allows fitting different SSMs within Bayesian hierarchical frameworks (61,88). Bayesian estimation allowed quantification of parameter estimates and uncertainty in the form of the posterior distribution. Before conducting any analyses on model parameters, we ensured model convergence by inspecting trace plots and using the Gelman-Rubin Ȓ statistic which was below the common threshold value of 1.1 for all parameters (89). To ensure that models fit the actual data, we also performed posterior predictive checks and computed quantile probability plots which allowed us to compare predicted versus actual data.

We used the default priors set in HDDM as explained elsewhere (61). Markov chain Monte-Carlo (MCMC) sampling methods were used to accurately approximate the posterior distributions of the estimated parameters. The models were run with three chains, and we sampled between 14,000 and 22,000 from the posterior (with burn-in between 10,000 and 18,000 samples) depending on whether trial-based neural activities were included as regressors (explained below). Statistical analyses were performed on the group mean posteriors following methods that have already been established in other reports (90–92). Specifically, Bayesian hypothesis testing was performed by analyzing the probability mass of the parameter region in question (estimated by the number of samples drawn from the posterior that fall in this region; for example, percentage of posterior samples greater than zero). We deemed parameters significant if 95% of the samples taken from their posterior probabilities were non-zero.

Comparing the performance of different SSM versions, we focused on the DDM (17) with and without across-trial variability parameters, as well as on the Ornstein-Uhlenbeck model with varying drift rate (93), and SSMs with (stochastic) relative evidence accumulation processes without across-trial variability parameters but with either linearly (94) or Weibull-informed collapsing boundaries (95). Aside from posterior predictive checks, the deviance information criterion (DIC) was used for model comparisons, where lower DIC values favor models with the highest likelihood and least number of parameters (96). Model specifications and comparison can be found in Supplementary Table 2. The best-fitting Weibull model captured responses and RTs for each condition with the approximated likelihood (based on the LAN extension of the HDDM toolbox) of the Wiener first passage time process (W) with Weibull- informed, time-dependent decision boundaries defined as:

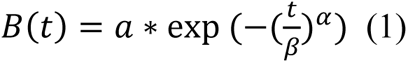

with *t* referring to (within-trial) time, *a* referring to boundary separation (i.e., the initial distance between the two decision thresholds at time 0 which indexes stimulus onset), *α* referring to the shape of the boundary collapse, and *β* referring to the onset of the boundary collapse. The Wiener process (W) is specified as:

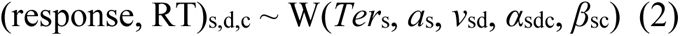

with *s* referring to subjects, *d* referring to discriminability (stronger, weaker), *c* referring to conflict (lower, higher), *v* referring to drift rate, and *Ter* referring to nondecision time. The starting point (*z*) was fixed to half the decision boundary (*a*). For all subjects, model parameters were specified by regression-based equations (with *I* referring to subject-specific intercepts) as follows:

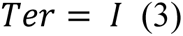

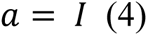

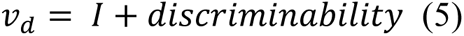

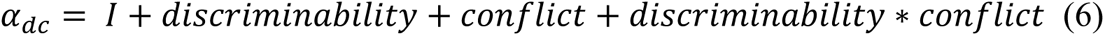

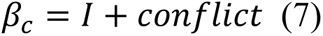

### Integrating electrophysiological data into the model-based analysis

After establishing that the Weibull DDM captured the behavioral patterns best, we augmented the model to determine whether trial-by-trial z-scored theta power (θz) influenced the decision threshold dynamics (reflected by the model parameters *α*, *β*) at the single trial level. Specifically, for quantifying theta activity during the pre-response period, we used z-scored mean activation during the last 500ms before response. For quantifying theta activity during the post-stimulus period, we used z-scored mean activation during the first 500ms after stimulus onset. As discussed in the Introduction, we focused on theta power based on strong a priori hypotheses established by previous work (34,42,49). Estimating the modulation of theta activity on decision boundary dynamics quantifies the psychological interpretation of these neural regressors. Thus, the regression-based equations for the shape (*α*) and rate (*β*) of the collapsing boundaries were augmented as follows:

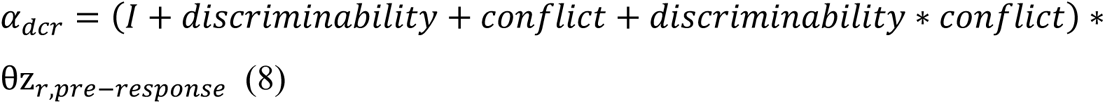

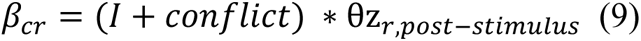

where *r* refers to trials, *θz* refers to the z-scored (hierarchically mean-centered) theta power during the pre-response period (i.e., time window between response and 500ms prior to response) and post-stimulus period (i.e., time window between stimulus onset and 500ms after stimulus onset). Equations for the other model parameters remained the same as for the initially established best-fitting Weibull model (see equations in previous subsection). We estimated the posteriors of coefficients for trial-specific regressors solely at the group level. This established approach allowed us to effectively handle possible collinearity among model parameters, stabilize parameter estimates, and avoid excessive parameter expansion (68,91). The determination of statistical significance for regression coefficients relied on the distribution of their posterior probabilities.

## Footnotes

1 Based on Wilcoxon signed-rank test: V = 316, N = 26, p < 0.001, r = -0.752. Supplementary Fig.1 provides details.

2 Based on Wilcoxon signed-rank test: V = 66, N = 26, p = 0.004, r = -0.560.

3 We did not find differences in drift rate or nondecision time between participants with recordings from the STN vs. the GP subsegments (Supplementary Table 4 & Fig. 3). Moreover, the students without neurological conditions showed similar modulations in the model parameters (Supplementary Fig. 4).

4 This is also reflected in the theta decline during the post-stimulus period shown in Fig. 3A (middle-left panel).

## References

1. Bogacz R, Gurney K. The Basal Ganglia and Cortex Implement Optimal Decision Making Between Alternative Actions. Neural Comput. 2007 Feb 1;19(2):442–77.

2. Drugowitsch J, Moreno-Bote R, Churchland AK, Shadlen MN, Pouget A. The Cost of Accumulating Evidence in Perceptual Decision Making. J Neurosci. 2012 Mar 14;32(11):3612–28.

3. Thura D, Cisek P. The Basal Ganglia Do Not Select Reach Targets but Control the Urgency of Commitment. Neuron. 2017 Aug 30;95(5):1160–1170.e5.

4. Malhotra G, Leslie DS, Ludwig CJH, Bogacz R. Overcoming Indecision by Changing the Decision Boundary. J Exp Psychol Gen. 2017 Jun;146(6):776–805.

5. Palestro JJ, Weichart E, Sederberg PB, Turner BM. Some task demands induce collapsing bounds: Evidence from a behavioral analysis. Psychon Bull Rev. 2018 Aug 1;25(4):1225– 48.

6. Shadlen MN, Newsome WT. Neural Basis of a Perceptual Decision in the Parietal Cortex (Area LIP) of the Rhesus Monkey. J Neurophysiol. 2001 Oct;86(4):1916–36.

7. Smith Y, Bevan MD, Shink E, Bolam JP. Microcircuitry of the direct and indirect pathways of the basal ganglia. Neuroscience. 1998 Sep 1;86(2):353–87.

8. Ding L, Gold JI. The Basal Ganglia’s Contributions to Perceptual Decision Making. Neuron. 2013 Aug;79(4):640–9.

9. Richard Ridderinkhof K, Forstmann BU, Wylie SA, Burle B, van den Wildenberg WPM. Neurocognitive mechanisms of action control: resisting the call of the Sirens. WIREs Cogn Sci. 2011;2(2):174–92.

10. Doi T, Fan Y, Gold JI, Ding L. The caudate nucleus contributes causally to decisions that balance reward and uncertain visual information. Lee D, Frank MJ, Lee D, Chandrasekaran C, editors. eLife. 2020 Jun 22;9:e56694.

11. Yartsev MM, Hanks TD, Yoon AM, Brody CD. Causal contribution and dynamical encoding in the striatum during evidence accumulation. Gold JI, Behrens TE, Gold JI, Humphries MD, editors. eLife. 2018 Aug 24;7:e34929.

12. Ding L, Gold JI. Caudate Encodes Multiple Computations for Perceptual Decisions. J Neurosci. 2010 Nov 24;30(47):15747–59.

13. Moss MM, Zatka-Haas P, Harris KD, Carandini M, Lak A. Dopamine Axons in Dorsal Striatum Encode Contralateral Visual Stimuli and Choices. J Neurosci. 2021 Aug 25;41(34):7197–205.

14. Westbrook A, Frank MJ, Cools R. A mosaic of cost–benefit control over cortico-striatal circuitry. Trends Cogn Sci. 2021 Aug 1;25(8):710–21.

15. Nambu A, Tokuno H, Hamada I, Kita H, Imanishi M, Akazawa T, et al. Excitatory Cortical Inputs to Pallidal Neurons Via the Subthalamic Nucleus in the Monkey. J Neurophysiol. 2000 Jul;84(1):289–300.

16. Gurney K, Prescott TJ, Redgrave P. A computational model of action selection in the basal ganglia. I. A new functional anatomy. Biol Cybern. 2001 May 1;84(6):401–10.

17. Ratcliff R. A theory of memory retrieval. Psychol Rev. 1978;85(2):59–108.

18. Wald A, Wolfowitz J. Bayes Solutions of Sequential Decision Problems. Proc Natl Acad Sci. 1949 Feb;35(2):99–102.

19. Ging-Jehli NR, Ratcliff R, Arnold LE. Improving neurocognitive testing using computational psychiatry—A systematic review for ADHD. Psychol Bull. 2021;147(2):169–231.

20. Smith PL, Ratcliff R. Psychology and neurobiology of simple decisions. Trends Neurosci. 2004 Mar 1;27(3):161–8.

21. Forstmann BU, Wagenmakers EJ. Model-Based Cognitive Neuroscience: A Conceptual Introduction. In: Forstmann BU, Wagenmakers EJ, editors. An Introduction to Model-Based Cognitive Neuroscience [Internet]. New York, NY: Springer; 2015 [cited 2022 Mar 24]. p. 139–56. Available from: 10.1007/978-1-4939-2236-9_7

22. Ratcliff R, Frank MJ. Reinforcement-Based Decision Making in Corticostriatal Circuits: Mutual Constraints by Neurocomputational and Diffusion Models. Neural Comput. 2012 May 1;24(5):1186–229.

23. Fengler A, Bera K, Pedersen ML, Frank MJ. Beyond Drift Diffusion Models: Fitting a Broad Class of Decision and Reinforcement Learning Models with HDDM. J Cogn Neurosci. 2022 Sep 1;34(10):1780–805.

24. Aron AR, Herz DM, Brown P, Forstmann BU, Zaghloul K. Frontosubthalamic Circuits for Control of Action and Cognition. J Neurosci. 2016 Nov 9;36(45):11489–95.

25. Wiecki TV, Frank MJ. A computational model of inhibitory control in frontal cortex and basal ganglia. Psychol Rev. 2013;120(2):329–55.

26. Zavala B, Zaghloul K, Brown P. The subthalamic nucleus, oscillations, and conflict. Mov Disord. 2015;30(3):328–38.

27. Frank MJ. Hold your horses: A dynamic computational role for the subthalamic nucleus in decision making. Neural Netw. 2006 Oct 1;19(8):1120–36.

28. Moolchand P, Jones SR, Frank MJ. Biophysical and Architectural Mechanisms of Subthalamic Theta under Response Conflict. J Neurosci. 2022 Jun 1;42(22):4470–87.

29. Frank MJ, Samanta J, Moustafa AA, Sherman SJ. Hold Your Horses: Impulsivity, Deep Brain Stimulation, and Medication in Parkinsonism. Science. 2007 Nov 23;318(5854):1309–12.

30. Schmidt R, Leventhal DK, Mallet N, Chen F, Berke JD. Canceling actions involves a race between basal ganglia pathways. Nat Neurosci. 2013 Aug;16(8):1118–24.

31. Eagle DM, Baunez C, Hutcheson DM, Lehmann O, Shah AP, Robbins TW. Stop-Signal Reaction-Time Task Performance: Role of Prefrontal Cortex and Subthalamic Nucleus. Cereb Cortex. 2008 Jan 1;18(1):178–88.

32. Herz DM, Zavala BA, Bogacz R, Brown P. Neural Correlates of Decision Thresholds in the Human Subthalamic Nucleus. Curr Biol. 2016 Apr;26(7):916–20.

33. Wessel JR, Waller DA, Greenlee JD. Non-selective inhibition of inappropriate motor-tendencies during response-conflict by a fronto-subthalamic mechanism. Verstynen T, Ivry RB, Verstynen T, Fountas Z, editors. eLife. 2019 May 7;8:e42959.

34. Herz DM, Tan H, Brittain JS, Fischer P, Cheeran B, Green AL, et al. Distinct mechanisms mediate speed-accuracy adjustments in cortico-subthalamic networks. Lakatos P, editor. eLife. 2017 Jan 31;6:e21481.

35. Zavala BA, Tan H, Little S, Ashkan K, Hariz M, Foltynie T, et al. Midline frontal cortex low-frequency activity drives subthalamic nucleus oscillations during conflict. J Neurosci Off J Soc Neurosci. 2014 May 21;34(21):7322–33.

36. Zavala B, Damera S, Dong JW, Lungu C, Brown P, Zaghloul KA. Human Subthalamic Nucleus Theta and Beta Oscillations Entrain Neuronal Firing During Sensorimotor Conflict. Cereb Cortex. 2017 Jan 1;27(1):496–508.

37. Isoda M, Hikosaka O. Role for Subthalamic Nucleus Neurons in Switching from Automatic to Controlled Eye Movement. J Neurosci. 2008 Jul 9;28(28):7209–18.

38. Yoshida M, Precht W. Monosynaptic inhibition of neurons of the substantia nigra by caudatonigral fibers. Brain Res. 1971 Sep 10;32(1):225–8.

39. Bauswein E, Fromm C, Preuss A. Corticostriatal cells in comparison with pyramidal tract neurons: contrasting properties in the behaving monkey. Brain Res. 1989 Jul 24;493(1):198–203.

40. Maanen L van, Brown SD, Eichele T, Wagenmakers EJ, Ho T, Serences J, et al. Neural Correlates of Trial-to-Trial Fluctuations in Response Caution. J Neurosci. 2011 Nov 30;31(48):17488–95.

41. Mansfield EL, Karayanidis F, Jamadar S, Heathcote A, Forstmann BU. Adjustments of Response Threshold during Task Switching: A Model-Based Functional Magnetic Resonance Imaging Study. J Neurosci. 2011 Oct 12;31(41):14688–92.

42. Cavanagh JF, Wiecki TV, Cohen MX, Figueroa CM, Samanta J, Sherman SJ, et al. Subthalamic nucleus stimulation reverses mediofrontal influence over decision threshold. Nat Neurosci. 2011 Nov;14(11):1462–7.

43. Zuure MB, Hinkley LB, Tiesinga PHE, Nagarajan SS, Cohen MX. Multiple Midfrontal Thetas Revealed by Source Separation of Simultaneous MEG and EEG. J Neurosci. 2020 Sep 30;40(40):7702–13.

44. Muralidharan V, Aron AR, Cohen MX, Schmidt R. Two modes of midfrontal theta suggest a role in conflict and error processing. NeuroImage. 2023 Jun 1;273:120107.

45. Zeng K, Drummond NM, Ghahremani A, Saha U, Kalia SK, Hodaie M, et al. Fronto-subthalamic phase synchronization and cross-frequency coupling during conflict processing. NeuroImage. 2021 Sep 1;238:118205.

46. Wei W, Rubin JE, Wang XJ. Role of the Indirect Pathway of the Basal Ganglia in Perceptual Decision Making. J Neurosci. 2015 Mar 4;35(9):4052–64.

47. Yoshida A, Tanaka M. Two Types of Neurons in the Primate Globus Pallidus External Segment Play Distinct Roles in Antisaccade Generation. Cereb Cortex. 2016 Mar 1;26(3):1187–99.

48. Dong J, Hawes S, Wu J, Le W, Cai H. Connectivity and Functionality of the Globus Pallidus Externa Under Normal Conditions and Parkinson’s Disease. Front Neural Circuits [Internet]. 2021 [cited 2023 Sep 7];15. Available from: https://www.frontiersin.org/articles/10.3389/fncir.2021.645287

49. Aron AR, Poldrack RA. Cortical and Subcortical Contributions to Stop Signal Response Inhibition: Role of the Subthalamic Nucleus. J Neurosci. 2006 Mar 1;26(9):2424–33.

50. Palmer J, Huk AC, Shadlen MN. The effect of stimulus strength on the speed and accuracy of a perceptual decision. J Vis. 2005 May 2;5(5):1.

51. Roitman JD, Shadlen MN. Response of Neurons in the Lateral Intraparietal Area during a Combined Visual Discrimination Reaction Time Task. J Neurosci. 2002 Nov 1;22(21):9475–89.

52. Ratcliff R, Cherian A, Segraves M. A Comparison of Macaque Behavior and Superior Colliculus Neuronal Activity to Predictions From Models of Two-Choice Decisions. J Neurophysiol. 2003 Sep;90(3):1392–407.

53. Redgrave P, Prescott TJ, Gurney K. The basal ganglia: A vertebrate solution to the selection problem? Neuroscience. 1999 Apr;89(4):1009–23.

54. Gold JI, Shadlen MN. Banburismus and the brain: decoding the relationship between sensory stimuli, decisions, and reward. Neuron. 2002 Oct 10;36(2):299–308.

55. Gold JI, Shadlen MN. The Influence of Behavioral Context on the Representation of a Perceptual Decision in Developing Oculomotor Commands. J Neurosci. 2003 Jan 15;23(2):632–51.

56. Jazayeri M, Movshon JA. Integration of sensory evidence in motion discrimination. J Vis. 2007 Sep 20;7(12):7.

57. Jazayeri M, Movshon JA. A new perceptual illusion reveals mechanisms of sensory decoding. Nature. 2007 Apr;446(7138):912–5.

58. Swaminathan SK, Freedman DJ. Preferential encoding of visual categories in parietal cortex compared with prefrontal cortex. Nat Neurosci. 2012 Feb;15(2):315–20.

59. Botvinick MM, Braver TS, Barch DM, Carter CS, Cohen JD. Conflict monitoring and cognitive control. Psychol Rev. 2001;108(3):624–52.

60. Botvinick M, Plaut DC. Representing task context: proposals based on a connectionist model of action. Psychol Res. 2002 Nov 1;66(4):298–311.

61. Wiecki T, Sofer I, Frank M. HDDM: Hierarchical Bayesian estimation of the Drift-Diffusion Model in Python. Front Neuroinformatics [Internet]. 2013 [cited 2022 Mar 24];7. Available from: https://www.frontiersin.org/article/10.3389/fninf.2013.00014

62. 62. Kruschke J. Doing Bayesian Data Analysis: A Tutorial with R, JAGS, and Stan. Academic Press; 2014. 772 p.

63. Herz DM, Little S, Pedrosa DJ, Tinkhauser G, Cheeran B, Foltynie T, et al. Mechanisms Underlying Decision-Making as Revealed by Deep-Brain Stimulation in Patients with Parkinson’s Disease. Curr Biol CB. 2018 Apr 23;28(8):1169–1178.e6.

64. Parent A, Hazrati LN. Functional anatomy of the basal ganglia. II. The place of subthalamic nucleus and external pallidium in basal ganglia circuitry. Brain Res Rev. 1995 Jan 1;20(1):128–54.

65. Stoessl AJ, Lehericy S, Strafella AP. Imaging insights into basal ganglia function, Parkinson’s disease, and dystonia. The Lancet. 2014 Aug 9;384(9942):532–44.

66. López-Azcárate J, Tainta M, Rodríguez-Oroz MC, Valencia M, González R, Guridi J, et al. Coupling between Beta and High-Frequency Activity in the Human Subthalamic Nucleus May Be a Pathophysiological Mechanism in Parkinson’s Disease. J Neurosci. 2010 May 12;30(19):6667–77.

67. Fischer P, Pogosyan A, Green AL, Aziz TZ, Hyam J, Foltynie T, et al. Beta synchrony in the cortico-basal ganglia network during regulation of force control on and off dopamine. Neurobiol Dis. 2019 Jul 1;127:253–63.

68. Frank MJ, Gagne C, Nyhus E, Masters S, Wiecki TV, Cavanagh JF, et al. fMRI and EEG Predictors of Dynamic Decision Parameters during Human Reinforcement Learning. J Neurosci. 2015 Jan 14;35(2):485–94.

69. Green N, Bogacz R, Huebl J, Beyer AK, Kühn AA, Heekeren HR. Reduction of Influence of Task Difficulty on Perceptual Decision Making by STN Deep Brain Stimulation. Curr Biol. 2013;17(23):1681–4.

70. Bolkan SS, Stone IR, Pinto L, Ashwood ZC, Iravedra Garcia JM, Herman AL, et al. Opponent control of behavior by dorsomedial striatal pathways depends on task demands and internal state. Nat Neurosci. 2022 Mar;25(3):345–57.

71. Hawkins GE, Forstmann BU, Wagenmakers EJ, Ratcliff R, Brown SD. Revisiting the Evidence for Collapsing Boundaries and Urgency Signals in Perceptual Decision-Making. J Neurosci. 2015 Feb 11;35(6):2476–84.

72. O’Connell RG, Kelly SP. Neurophysiology of Human Perceptual Decision-Making. Annu Rev Neurosci. 2021 Jul 8;44(Volume 44, 2021):495–516.

73. Kelly SP, Corbett EA, O’Connell RG. Neurocomputational mechanisms of prior-informed perceptual decision-making in humans. Nat Hum Behav. 2021 Apr;5(4):467–81.

74. Chen J, Wang Q, Li N, Huang S, Li M, Cai J, et al. Dyskinesia is Closely Associated with Synchronization of Theta Oscillatory Activity Between the Substantia Nigra Pars Reticulata and Motor Cortex in the Off L-dopa State in Rats. Neurosci Bull. 2021 Mar;37(3):323–38.

75. Shine JM, Handojoseno AMA, Nguyen TN, Tran Y, Naismith SL, Nguyen H, et al. Abnormal patterns of theta frequency oscillations during the temporal evolution of freezing of gait in Parkinson’s disease. Clin Neurophysiol. 2014 Mar 1;125(3):569–76.

76. Sakellariou DF, Dall’Orso S, Burdet E, Lin JP, Richardson MP, McClelland VM. Abnormal microscale neuronal connectivity triggered by a proprioceptive stimulus in dystonia. Sci Rep. 2020 Nov 27;10(1):20758.

77. Mirzaei A, Kumar A, Leventhal D, Mallet N, Aertsen A, Berke J, et al. Sensorimotor Processing in the Basal Ganglia Leads to Transient Beta Oscillations during Behavior. J Neurosci. 2017 Nov 15;37(46):11220–32.

78. Diesburg DA, Greenlee JD, Wessel JR. Cortico-subcortical β burst dynamics underlying movement cancellation in humans. Swann NC, Ivry RB, Muralidharan V, Schmidt R, editors. eLife. 2021 Dec 7;10:e70270.

79. Navid MS, Kammermeier S, Niazi IK, Sharma VD, Vuong SM, Bötzel K, et al. Cognitive task-related oscillations in human internal globus pallidus and subthalamic nucleus. Behav Brain Res. 2022 Apr 29;424:113787.

80. Balogh L, Pulay A, Angyal N, Vincze K, Kilencz T, Nemoda Z, et al. Upside down: dissecting impulsivity in attention-deficit hyperactivity disorder through genotype-phenotype association analyses. Eur Psychiatry. 2022 Jun;65(S1):S227–8.

81. Kuntsi J, Pinto R, Price TS, van der Meere JJ, Frazier-Wood AC, Asherson P. The Separation of ADHD Inattention and Hyperactivity-Impulsivity Symptoms: Pathways from Genetic Effects to Cognitive Impairments and Symptoms. J Abnorm Child Psychol. 2014 Jan 1;42(1):127–36.

82. Ging-Jehli NR, Arnold LE, Roley-Roberts ME, deBeus R. Characterizing Underlying Cognitive Components of ADHD Presentations and Co-morbid Diagnoses: A Diffusion Decision Model Analysis. J Atten Disord. 2022 Mar 1;26(5):706–22.

83. Ging-Jehli NR, Ratcliff R. Effects of aging in a task-switch paradigm with the diffusion decision model. Psychol Aging. 2020;35(6):850–65.

84. Schmitz F, Krämer RJ. Task Switching: On the Relation of Cognitive Flexibility with Cognitive Capacity. J Intell. 2023 Apr;11(4):68.

85. Schmitz F, Voss A. Decomposing task-switching costs with the diffusion model. J Exp Psychol Hum Percept Perform. 2012;38(1):222–50.

86. Asaad WF, Santhanam N, McClellan S, Freedman DJ. High-performance execution of psychophysical tasks with complex visual stimuli in MATLAB. J Neurophysiol. 2013 Jan;109(1):249–60.

87. R Core Team. R: A language and environment for statistical computing [Internet]. Vienna, Austria: R Foundation for Statistical Computing; 2021. Available from: https://www.R-project.org/

88. Fengler A, Govindarajan LN, Chen T, Frank MJ. Likelihood approximation networks (LANs) for fast inference of simulation models in cognitive neuroscience. Wyart V, Behrens TE, Acerbi L, Daunizeau J, editors. eLife. 2021 Apr 6;10:e65074.

89. Gelman A, Rubin DB. Inference from Iterative Simulation Using Multiple Sequences. Stat Sci. 1992;7(4):457–72.

90. Cavanagh JF, Wiecki TV, Kochar A, Frank MJ. Eye tracking and pupillometry are indicators of dissociable latent decision processes. J Exp Psychol Gen. 2014;143(4):1476– 88.

91. Wiecki T, Sofer I, Frank M. HDDM: Hierarchical Bayesian estimation of the Drift-Diffusion Model in Python. Front Neuroinformatics [Internet]. 2013 [cited 2023 Sep 21];7. Available from: https://www.frontiersin.org/articles/10.3389/fninf.2013.00014

92. Pedersen ML, Ironside M, Amemori K ichi, McGrath CL, Kang MS, Graybiel AM, et al. Computational phenotyping of brain-behavior dynamics underlying approach-avoidance conflict in major depressive disorder. PLOS Comput Biol. 2021 May 10;17(5):e1008955.

93. Busemeyer JR, Townsend JT. Fundamental derivations from decision field theory. Math Soc Sci. 1992 Jun 1;23(3):255–82.

94. Forstmann BU, Ratcliff R, Wagenmakers EJ. Sequential Sampling Models in Cognitive Neuroscience: Advantages, Applications, and Extensions. Annu Rev Psychol. 2016 Jan 4;67(1):641–66.

95. Churchland AK, Kiani R, Shadlen MN. Decision-making with multiple alternatives. Nat Neurosci. 2008 Jun;11(6):693–702.

96. Gelman A, Hill J. Data Analysis Using Regression and Multilevel/Hierarchical Models. Cambridge University Press; 2006. 651 p.

